# Superinfection Exclusion of Alphaherpesviruses Interferes with Virion Trafficking

**DOI:** 10.1101/2021.12.16.473075

**Authors:** James P. Cwick, Jonathan I. Owen, Irina Kochetkova, Nick Van Horssen, Matthew P. Taylor

**Author notes:** Corresponding Author: Matthew P. Taylor.

## Abstract

Superinfection exclusion (SIE) is a phenomenon in which a primary viral infection interferes with secondary viral infections within that same cell. Although SIE has been observed across many viruses, it has remained relatively understudied. A recently characterized glycoprotein D (gD) -independent SIE of alphaherpesviruses presents a novel mechanism of co-infection restriction for Herpes Simplex Virus Type 1 (HSV-1) and Pseudorabies virus (PRV). In this study, we evaluated the role of multiplicity of infection (MOI), receptor expression, and trafficking of virions to gain greater insight into potential mechanisms of alphaherpesvirus SIE.

We observed that high MOI secondary viral infections were able to overcome SIE in a manner that was independent of receptor availability. Utilizing recombinant viruses expressing fluorescent protein fusions, we assessed virion localization during SIE through live fluorescent microscopy of dual-labeled virions and localization of capsid assemblies. Analysis of these assemblies confirmed changes in the distribution of capsids during SIE. These results indicate that SIE during PRV infection inhibits viral entry or fusion while HSV-1 SIE inhibits infection through a post-entry mechanism. Although the timing and phenotype of SIE is similar between alphaherpesviruses, the related viruses implement different mechanisms to restrict coinfection.

**IMPORTANCE:** Most viruses utilize a form of superinfection exclusion to conserve resources and control population dynamics. gD-dependent superinfection exclusion in alphaherpesviruses is well-documented. However, the under-characterized gD-independent SIE provides new insight into how alphaherpesviruses limit sequential infection. The observations described here demonstrate that gD-independent SIE differs between PRV and HSV-1. Comparing these differences provide new insights into the underlying mechanisms of SIE implemented by two related viruses.

## INTRODUCTION

Superinfection exclusion (SIE) is a virally induced process observed after a primary virus establishes infection in a cell which then prevents subsequent infection by a secondary virus (1,2). SIE serves as a strong, selective pressure to control competition for cellular resources and regulate viral population diversity (3–5). SIE has been observed across diverse viral families and differs in how it mechanistically alters secondary infection (6–10). For example, poxviruses employ multiple mechanisms to disrupt entry or fusion of superinfecting virions (7,11). In contrast, SIE implemented by different flaviviruses blocks viral RNA synthesis of superinfecting viral genomes, though mutations in specific structural proteins can circumvent this SIE mechanism (8). Mechanisms of SIE are ubiquitous across viral families and target every step of viral infection or replication (1). Important to this work, SIE has also been observed for alphaherpesviruses including Herpes Simplex virus type 1 (HSV-1) and Pseudorabies virus (PRV) (4,12,13).

Alphaherpesviruses are a family of neurotropic viruses with the ability to establish latency in neurons and have a broad impact on human health and agriculture. HSV-1 has achieved a global infection rate of 80% with approximately 40% of individuals seropositive within the United States (14,15). Documented cases of both symptomatic and asymptomatic shedding within infected individuals promote high circulating populations of HSV and the probability for sequential exposure (16–19). These conditions of viral prevalence and transmission increase the probability of superinfection within an individual that can lead to the development of recombinant genomes with novel phenotypes (20–22). Recombinant viruses generated from superinfection have been observed in many alphaherpesviruses including Varicella Zoster virus (VZV) and between HSV-1 and 2 (23–27). PRV greatly impacts agricultural practices and production. An estimated $500 million is spent annually in the United States for vaccination, disease surveillance, and livestock eradication (24). The prevalent use of glycoprotein-E negative vaccines has allowed possible recombination with circulating strains of PRV (28). Despite concerns about novel recombinant genomes, we still do not know much about the mechanisms that HSV-1 and PRV employ to interfere with superinfecting virions and limit co-infection.

Established literature of SIE for alphaherpesviruses have focused on the role and effect of glycoprotein D expression (gD) (29,30). However, another mechanism for alphaherpesvirus SIE has been observed that is independent of gD expression (4). This gD-independent SIE is potentially a departure from the canonical mechanism of occlusion or internalization of cellular receptor proteins like nectin-1 (31,32). Additionally, gD-independent SIE was dependent upon active viral replication and protein synthesis during the first 2 hours of primary viral infection to exclude secondary viral inoculum. However, these results could neither isolate the step of viral infection inhibited nor the mechanism contributing to gD-independent SIE (4). To better characterize and understand SIE for both HSV-1 and PRV, we applied a combination of live fluorescent microscopy with fluorescent protein expressing recombinant viruses to understand inhibition of secondary infection. We investigated the roles of multiplicity of infection (MOI), receptor internalization, and virion distribution to understand PRV and HSV-1 SIE. Together, results from this work will demonstrate that SIE interferes with early steps of viral infection to suppress the replication of superinfecting viruses.

## MATERIALS AND METHODS

### Cells and viruses

Porcine kidney epithelial cells (PK15), human keratinocyte cells (HaCat), and green African kidney (Vero) cell lines were maintained in Dulbecco’s modified Eagle’s medium (DMEM) supplemented with 10% (vol/vol) fetal bovine serum (FBS) and 1% (vol/vol) penicillin-streptomycin. PRV stocks and plaque assays were performed on PK15 cells as previously described (33). PRV isolates containing glycoprotein D deletion were propagated on PK15 cells that stably express glycoprotein D (g5 cells) needed for supporting virion infectivity (4,13,34). These cells were cultured as above but supplemented with histidinol (50 μg/mL) to maintain glycoprotein D expression. HSV stocks were propagated and performed on plaque assays with either HaCaT or Vero Cells (35).

All fluorescent PRV recombinants were derived from a PRV Becker background. PRV 287 and 289 express either an enhanced yellow fluorescent protein (eYFP) or cyan fluorescent protein (Turq2) fused to a 3xNLS signal peptide as described previously (4,34). PRV 180 contains an mRFP fusion to the minor capsid protein VP26 as previously characterized (36). PRV 137 expresses both an eGFP in-frame fusion to gM and an mRFP fusion to VP26 as previously described (37). PRV GS442 is a kind gift from Dr. Greg Smith and the Enquist Lab. PRV GS442 is a viral mutant lacking glycoprotein D expression. A PRV BAC was utilized to excise the gD gene via homologous recombination inserting a diffusible green fluorescent protein (eGFP). Originally, stocks of PRV GS442 were maintained in a mixture of G5 and PK15 cells in a ratio of approximately 3:1. Viral titers of PRV GS442 were determined with PK15 cells (4,13,34). Fluorescent HSV-1 recombinants were derived from HSV-1 strain 17. HSV-1 OK14 expresses an mRFP fusion to the capsid protein VP26 (36).

All stocks of fluorescent viruses were confirmed by fluorescence microscopy of plaques to ensure maintenance and fidelity of the associated fluorescent profiles. Experiments requiring fluorescent virions utilized fresh viral supernatants. Stocks were initiated with an MOI 10 infection of a 10 cm dish of support cells. At 24 hours post infection, media was collected and subjected to low-speed centrifugation to remove cells and debris. Media was supplemented with 20mM HEPES pH 7.0. Cell-free viral stocks of PRV 180 and HSV-1 OK14 were stored in 500 μL aliquots to maximize virion associated fluorescence.

### Microscopy

Images were acquired on a Nikon Ti-Eclipse (Nikon Instruments, Melville, NY) inverted microscope equipped with a SpectraX LED (Lumencor, Beaverton, OR) excitation module and fast-switching emission filter wheels (Prior Scientific, Rockland, MA). Fluorescence imaging used paired excitation/emission filters and dichroic mirrors for cyan fluorescent protein (CFP), YFP, and RFP (Chroma Technology Corp., Bellow Falls, VT). Brightfield images utilized a Plan Fluor 20 phase contrast (Ph) objective and an iXon 896 EM-CCD (Andor Technology Ltd., Belfast, Northern Ireland) camera using NIS Elements software. Acquisition of fluorescent virions utilized a CFI Plan Apochromat Lambda 60x and 100x oil immersion objective to visualize individual virion localization within infected cells. Infected cells were imaged at the time post-infection indicated in each figure legend, dependent on experimental design. Experiments were tested at least twice with multiple fields acquired for each condition of the experiment. Live imaging was performed on the microscope in conjunction with a stage-top incubation system [Quorum Scientific, Puslinch, ON, CA]. PK15 cells were cultured on Lab-Tek II chambered coverglass. Prior to imaging, a nonfluorescent primary virus was used to establish primary infection, washed, and then feed with media for 2 hours before the secondary infection was applied. Cells were maintained at 37 °C in a 5% (vol/vol) CO2 enriched atmosphere using a stage top incubator system. Imaging began directly after application of fluorescent viral inoculum, as described. Images were acquired every 2.5 minutes and fluorescent illumination intensity was set to 35% power with less than 100 milliseconds exposure for each channel to avoid photobleaching.

### Flow cytometry

Detection of extracellular nectin-1 in infected cells was performed using a BD LSR II or BD LSR Fortessa cytometer (BD Biosciences, San Jose, CA). Infected cells were harvested by 5mM EDTA supplemented with Dulbecco’s PBS (D-PBS) and washed once with Ca2+ and Mg2+ -free D-PBS with 0.1% BSA (flow cytometry buffer). Cells were then fixed 2% PFA, washed with 1x PBS solution, and resuspended in 200 μl of flow cytometry buffer. Fixed cells were incubated with mouse IgG monoclonal nectin-1-specific antibody (CK8) [ThermoFisher, Waltham, MA] at 2.5 μg/ml for 1 hour at room temperature. Cells were washed again and subsequently incubated with secondary goat anti-mouse IgG conjugated to Alexa Fluor 488 [ThermoFisher, Waltham, MA]. After 1 hour of incubation, cells were washed with 1x PBS solution and resuspended in flow cytometry buffer for analysis. Acquisition and gating were set on mock-infected and non-specific, secondary antibody-stained infected cell populations. All cytometry data were analyzed using the FlowJo data analysis software (FlowJo, LLC, Ashland, OR).

### MOI dependency for superinfected cells

FP expression in infected cells was assessed through fluorescent microscopy. The day before experiments, 1.0×10^5^ PK15 cells per well were seeded in 12-well plates. The next day, at time T=-1 hours, initial inoculations across four MOI conditions (10, 25, 50, and 100) were performed with PRV 289. One-hour post-inoculation, inoculum was aspirated, and cells were washed with 2 mL PBS per well. Then, 1 mL of viral media (DMEM, 2% FBS, 1% Pen-Strep) was supplemented into each well. This time point, T=0 hours, was taken to be the time of initial infection. At T=2 hours, media was aspirated, cells were again washed with 2 mL PBS per well, and secondary inoculations were performed with PRV 287. These infections were also performed across four MOI conditions (10, 25, 50, and 100), such that a 4×4 matrix was generated whereby every initial infection condition was subjected to every superinfection condition. At T=3 hours, superinfecting inoculum was aspirated, cells were washed with PBS, and cells were again overlaid with 1 mL viral media per well. Finally, at T=6 hours, cells were imaged on brightfield, CFP, and YFP channels using paired fluorescent filters. In certain experiments, cycloheximide (CHX) was used to inhibit protein synthesis in the initial inoculation. In these cases, after the wash of the initial inoculum at T=0, cells were overlaid with 1 mL viral media per well supplemented with 100 μg/mL cycloheximide (CHX). Consequently, at T=2 hours cells were subjected to two washes with PBS prior to application of the second inoculum, to ensure removal of any CHX during subsequent incubations.

### Virion localization

Fluorescent virion assemblies were imaged on an epifluorescent microscope and photomicrographs were acquired with 60x and 100x magnification objectives. Sequential photos of infected cells through the z-axis (z-stacks) were acquired and used to generate a 3-D reconstruction with NIS-Elements software to facilitate analysis of capsid localization on a per cell basis. Individual capsid assemblies were then counted in a single-cell analysis based upon their localization in the cell: perinuclear (adjacent to nuclear region of cell), intracellular virions (inside cellular membrane), and external virions (virions attached but unentered). Average counts of virions were then compared across all conditions and represented out of total virions.

The following conditions were used for experimental procedures: direct infection (DI), SIE, and SIE with CHX. Direct infections were established by the inoculation of mRFP-VP26 virions (HSV-1 OK14 and PRV 180) onto Vero or PK15 cells, respectively, for a 1-hour duration in a 37°C incubator. For SIE, cells were initially inoculated for 1 hour at 37°C with a non-fluorescent HSV-1 or PRV wild-type virus. Cells were then washed and maintained for 3 hours at 37°C. Afterward, they were subsequently infected with mRFP-VP26 labeled virions as described for direct infection. For SIE with CHX, cells were initially inoculated for 1 hour at 37°C with a non-fluorescent HSV-1 or PRV wild-type virus, then washed and supplemented with media containing CHX at 100 μg/mL during the 3-hour incubation at 37°C. After incubation, cells were subsequently infected with mRFP-VP26 labeled virions as described for direct infection. One hour after application of fluorescent virions, cells were washed and supplemented with fresh cellular media.

### Statistical analysis of virion localization

To compare the statistical differences between conditions of the virion localization experiments, statistical analysis was performed through GraphPad Prism (GraphPad Software, San Diego, California USA). To process the data derived from capsid localization, descriptive statistics was used in Prism to determine basic statistical parameters of quartiles, mean, and 95% confidence interval. D’Agostino & Pearson tests were applied to confirm that the data sets did not follow parameters of normal standardization due to the presence of outliers. Statistical comparisons were done with Kruskal Wallis tests, with Dunn’s multiple comparisons test as a means for comparing differences between conditions (SIE, DI, and SIE with CHX). One-Way ANOVA graphs depicting statistical significance and single cell analysis comparing the localization of viral capsids among conditions are shown in Figures 4–6.

## RESULTS

### PRV SIE is dependent upon MOI

SIE is a virally induced process that can directly or indirectly inhibit or block the course of secondary virion infection (1). However, the timing and extent of SIE for any virus is dependent on several variables of infection, including the inoculating dose. The number of virions infecting a single cell, the multiplicity of infection (MOI), has an immense effect on outcomes of viral replication. In previously published data, we established the timing of HSV-1 and PRV SIE under equivalent inoculation conditions of MOI 10. We hypothesized that SIE established by the initial MOI of the primary virus might be overcome with varying MOIs of the secondary virus.

To test the effect of MOI on SIE, we evaluated the effect of different doses for the initial and superinfecting inoculations. For all infections, application of the secondary viral inoculum occurred 2 hours after removal of the primary inoculum. To detect exclusion of the secondary inoculum, we evaluated fluorescent protein expression from recombinant viruses. Fluorescent protein expression correlating with either primary (CFP) or secondary (YFP) infection was evaluated by fluorescent microscopy at 7-9 hours post inoculation (Figure 1a). The extent of YFP expression in the population of infected cells inversely reflects the degree of SIE. Fewer YFP positive nuclei indicate greater SIE, while more YFP positive nuclei indicate less SIE. In line with our previous report, we observe extensive SIE when primary and secondary inoculations are an equal MOI (Figure 1b). Conditions where the secondary MOI was lower than the initial infection resulted in reduced YFP positive nuclei, indicative of greater or more effective SIE. In line with our hypothesis, higher MOIs for the secondary inoculation were able to overcome SIE, resulting in greater numbers of YFP expressing cells. The greatest difference of YFP expression occurred with a low primary inoculation and a high secondary MOI (MOI 10 primary, 100 secondary). The extent of FP expression under these conditions was nearly equivalent to that observed with simultaneous inoculation of the two viruses, suggesting no exclusion of the secondary virus.

**Figure 1.**
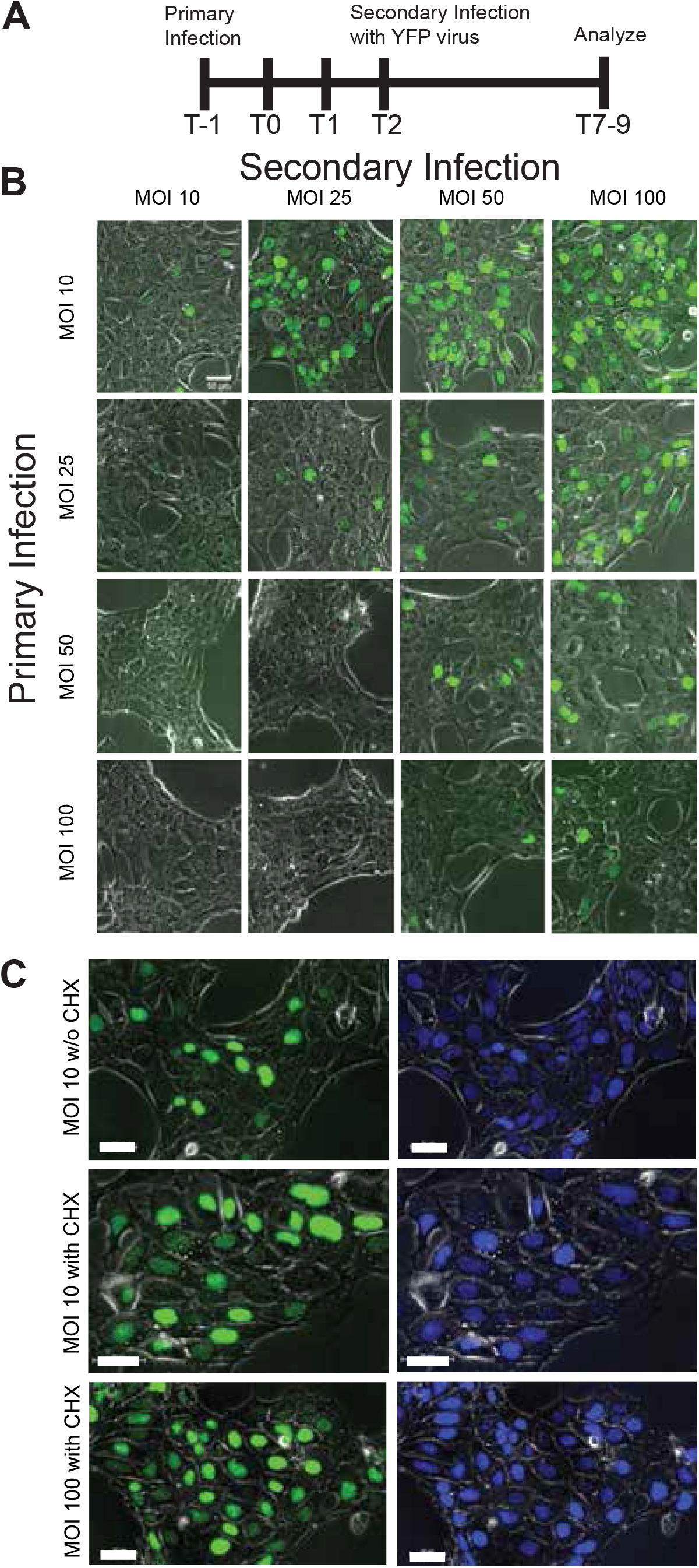
SIE dependence on primary and secondary MOIs. Experiments done in triplicate. A) Diagram of the timeline of experimentation. B) Comparison of SIE across a 4×4 matrix of primary and secondary infections. Top row corresponds to the MOI of the secondary virus (YFP labeled), while the vertical column corresponds to the MOI of the primary virus. Images were acquired and analyzed 7-9 hours after initial infection (or 4-6 hours after superinfection). C) Analysis of SIE with CHX treatment. Experiments were repeated with previously described SIE conditions. CHX treatment occurred between the application of the primary and secondary inoculum. Top row depicts fluorescent protein expression without CHX (MOI 10), while the middle (MOI 10) and the bottom (MOI 100) depict fluorescent protein expression with CHX treatment.

In previous work, it was observed that alphaherpesvirus SIE could be disrupted using the protein synthesis inhibitor cycloheximide (CHX) (4). To determine if protein synthesis is necessary for SIE at the different inoculating doses, we utilized CHX treatment to alleviate SIE at the extremes of our inoculating doses of MOI 10 and 100. Fluorescent protein detection during SIE was repeated with CHX treatment during the 2-hour window between primary and secondary inoculation. CHX treatment during both MOI 10 and 100 infections alleviated SIE, with an increase of YFP expressing cells (Figure 1c). Importantly, the secondary virus appears to exhibit relatively similar fluorescence profiles at both conditions, consistent with a similar mechanism of exclusion independent of MOI.

The data supports that PRV SIE is dependent upon MOI of both the primary and secondary virus. Alleviation of SIE occurred by either an overwhelming inoculating dose of secondary virus or through inhibition of protein synthesis by CHX treatment. In both cases, protein production by either cellular or viral origins still plays an important role in the dynamics contributing to SIE establishment.

### SIE is independent of surface receptor modulation

Since other forms of SIE disrupt receptor-mediated entry of virions, a possible explanation for SIE disruption of alphaherpesviruses could be dependent upon modulation of available surface receptors. Indeed, HSV-1 infection has been previously shown to internalize surface expression of cellular entry receptors like nectin-1 (39,40). The expected modulation of cellular entry receptors could affect SIE by decreasing the availability of cellular receptors for attaching virions, thus decreasing entry in a manner independent of gD expression. Based on these reports, we hypothesized that nectin-1 surface expression changes contribute to SIE.

To address this hypothesis, nectin-1 surface expression was evaluated during PRV infection. Changes in surface nectin-1 detection was correlated to the time of SIE induction following primary infection. PK15 cells were either mock-infected or infected with wild-type PRV. Following infection, cells were then either mock treated or treated with CHX. At the indicated times post-infection, cells were fixed, stained, and assessed for surface nectin-1 through mean-fluorescence intensity by flow cytometry.

We focused our analysis at 0-, 1-, and 2-hours post-inoculum removal to evaluate cell surface nectin during the time of SIE induction. For T0, there was a significant drop in the surface of nectin-1 expression during PRV infection. This outcome supports past literature that observable internalization of nectin-1 occurs following alphaherpesvirus infection (38,39). However, T1 observes no notable difference regarding surface expression of nectin-1 for PRV infection with or without CHX treatment (Figure 2). Overall, the data demonstrates that relative expression of nectin-1 for CHX treated and untreated conditions remains the same. Although CHX inhibits SIE, it does reduce surface nectin-1 levels. This supports a model that reduced nectin-1 expression following primary virion entry is still sufficient for effective entry of a secondary virus. These results surmise that any potential SIE mechanism is not mediated by changes in nectin-1 surface expression.

**Figure 2.**
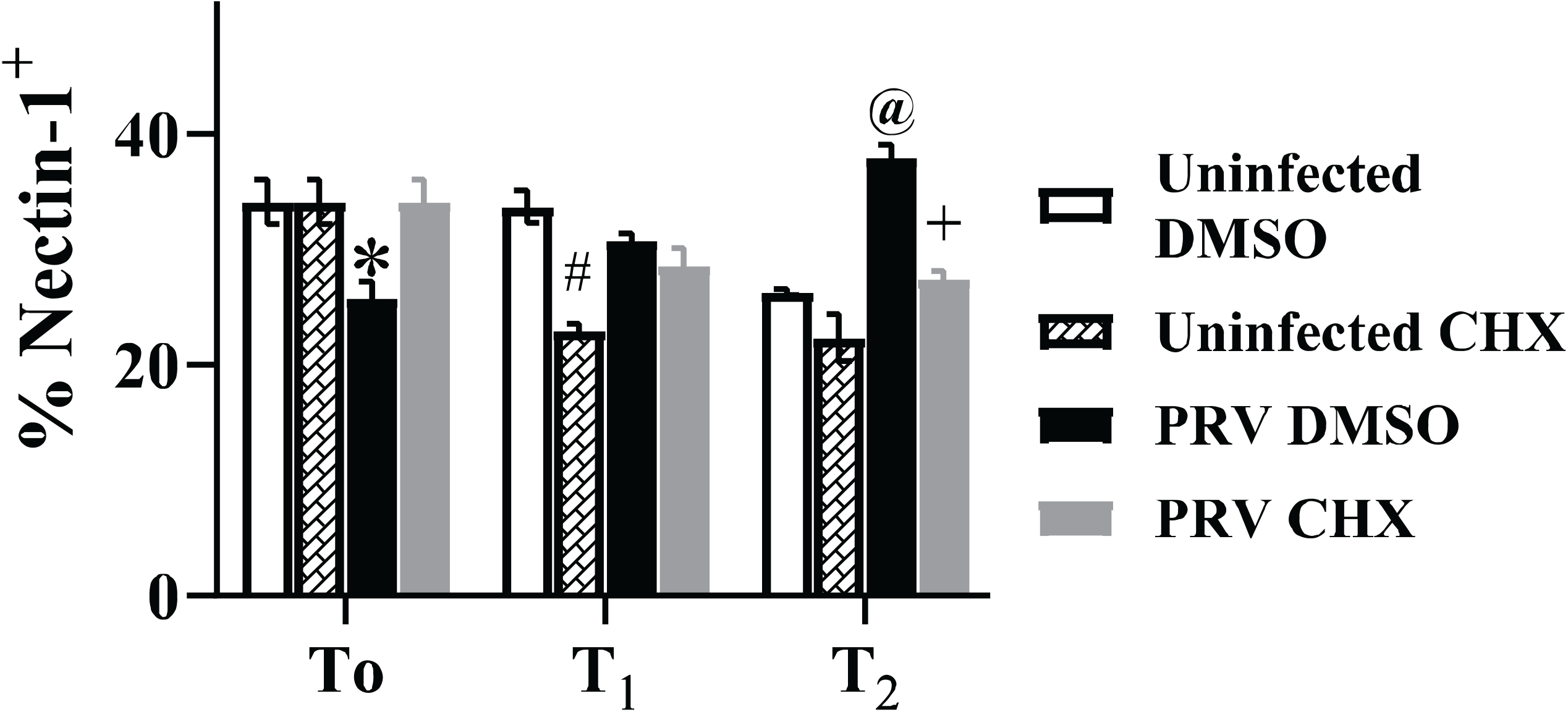
Cell surface nectin expression does not correlate with SIE during PRV infection. Depicted is the percentage of nectin-1 expression on the surface of the cells during PRV infection (MOI 50) at hours after inoculum removal. Cells were gated for being positive of stained fluorescence as compared to unstained controls. * represents a p-value of <0.05 as compared to uninfected DMSO. # represents a p value of <0.005 as compared to uninfected DMSO control. @ represents a p value of <0.001 as compared to uninfected DMSO control. + represented a p value of <0.005 as compared to PRV DMSO.

### SIE impacts entry of secondary virions

Our previous experiments using fluorescent viruses indicate that glycoprotein D independent SIE interferes with secondary viral infection. However, it could not isolate the step(s) of infection that was inhibited (4). Inhibition of one or several steps of infection–including attachment, entry, capsid trafficking, or transcription–could result in inhibition of secondary virus associated FP expression (41,42). To understand the extent of secondary virion inhibition during SIE, we developed a method to monitor virion entry and trafficking using direct FP fusions to virion components.

To determine how SIE impacts early steps of viral entry, time-lapse microscopy was performed with a dual fluorescently labeled virion (PRV 137). As modeled in Figure 3a, PRV137 produces virions that incorporate a GFP fusion to viral protein gM and a mRFP fusion to the capsid associated protein VP26 (38). Colocalization of both fluorophores indicates an intact virion, while dissociation of the two fluorescent signals correlates with entry of the capsid to the cell. The gM protein should remain at the plasma membrane, while the VP26 viral capsid protein will traffic towards the nucleus for insertion of the viral genome and subsequent viral replication. Microscopic examination of secreted PRV 137 virions used for experimental inoculation observed approximately 75% of fluorescent capsid assemblies (red puncta) contain detectable amounts of GFP (Figure 3b). Therefore, these co-labeled virions provide an excellent means to evaluate the distribution and structure of virions during entry.

**Figure 3.**
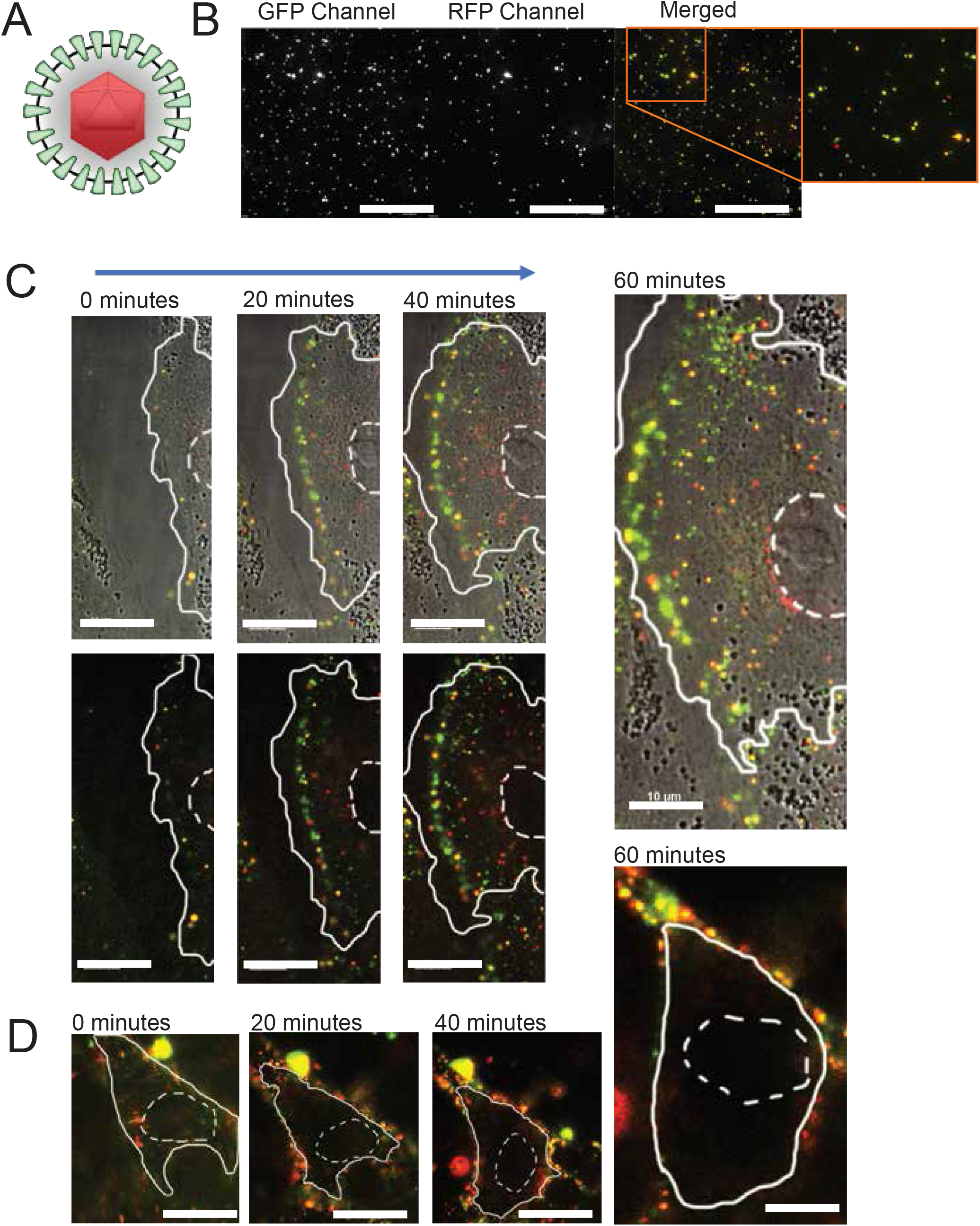
Live fluorescent microscopy observes absence of nuclear localized capsids during SIE. A) Model of PRV 137 virion containing a mRFP-VP26 fusion (red capsid) and a gM-eGFP fusion (green envelope protein). B) Distribution of PRV 137 fluorescent assemblies from a viral supernatant preparation. Images split into designated fluorescent channels and composite image. C) Select images from a time course of direct infection with PRV 137. Images taken every 20 minutes. Top row corresponds to brightfield and fluorescent composite images, while the bottom row demonstrates only fluorescent merged images. Nucleus is outlined with dashed lines while the cell boundary is outlined with a solid line. D) Select images from a time course of PRV 137 during SIE. Images were taken every 20 minutes after application of PRV 137. Cell boundaries as depicted above.

To evaluate virion entry under SIE conditions, sequential micrographs of cells were acquired using a live-imaging approach after inoculation (37). For our live-imaging experiment, we evaluated virion localization and fluorescence during conditions of direct infection (DI) and SIE. For DI, PRV 137 at an MOI 100 was applied to uninfected cells. In parallel for SIE, PRV 137 was applied to cells that had been previously infected with a non-fluorescent PRV 2-hours prior. Image acquisition for both conditions began immediately after application of the PRV 137 inoculum and every 2.5 minutes for a period of one and a half hours.

Representative images at 20 minutes intervals from the resulting time-lapse movies (Supplemental Movies 1 and 2) are depicted for both DI and SIE. Under conditions of DI (Figure 3c), we observed separation of fluorescence, with an increased number of red punctate structures within the cell. Capsid internalization and subsequent trafficking towards the nucleus are evidenced by red puncta near the nucleus. In contrast, cells superinfected with PRV 137 (SIE condition) exhibit no internalized or nuclear-associated capsid assemblies (Figure 3d). In fact, most of the fluorescent structures remain at or near the cellular boundary.

These observations are consistent with our expectations that internalization or trafficking of secondary virions is strongly suppressed during SIE. The lack of separation between the two fluorescent labels suggests an impairment of virion fusion, while the absence of internalized red fluorescent capsids is evidence of impaired capsid trafficking. To further evaluate the impact of SIE on secondary virion entry, we sought to develop more quantitative approaches to understand virion trafficking.

### SIE in PRV reduces capsid entry

To quantify the extent of virion entry and trafficking, we evaluated the distribution of fluorescent capsids within a 3-dimensional reconstruction of infected cells. Cells were inoculated with mRFP-VP26 labeled capsids (PRV 180 or HSV-1 OK14) under conditions of direct infection, SIE, and SIE with treatment of cycloheximide (CHX). Following infection, cells were sequentially imaged through their z-axis to acquire a full imaging volume of the cells. After image acquisition, cells were rendered into 3-D models and individual fluorescent puncta were counted and categorized by relative location in each cell: capsids near the nuclear region (nuclear), within the cellular boundary (intracellular), and capsids outside of the cellular boundary (peripheral). Analysis of infected cells from each condition excluded cells with extensive fluorescence aggregation or those that contained overlapping cellular boundaries. Representative images of infections under the different conditions are depicted in Figure 4, 5 and 6.

**Figure 4.**
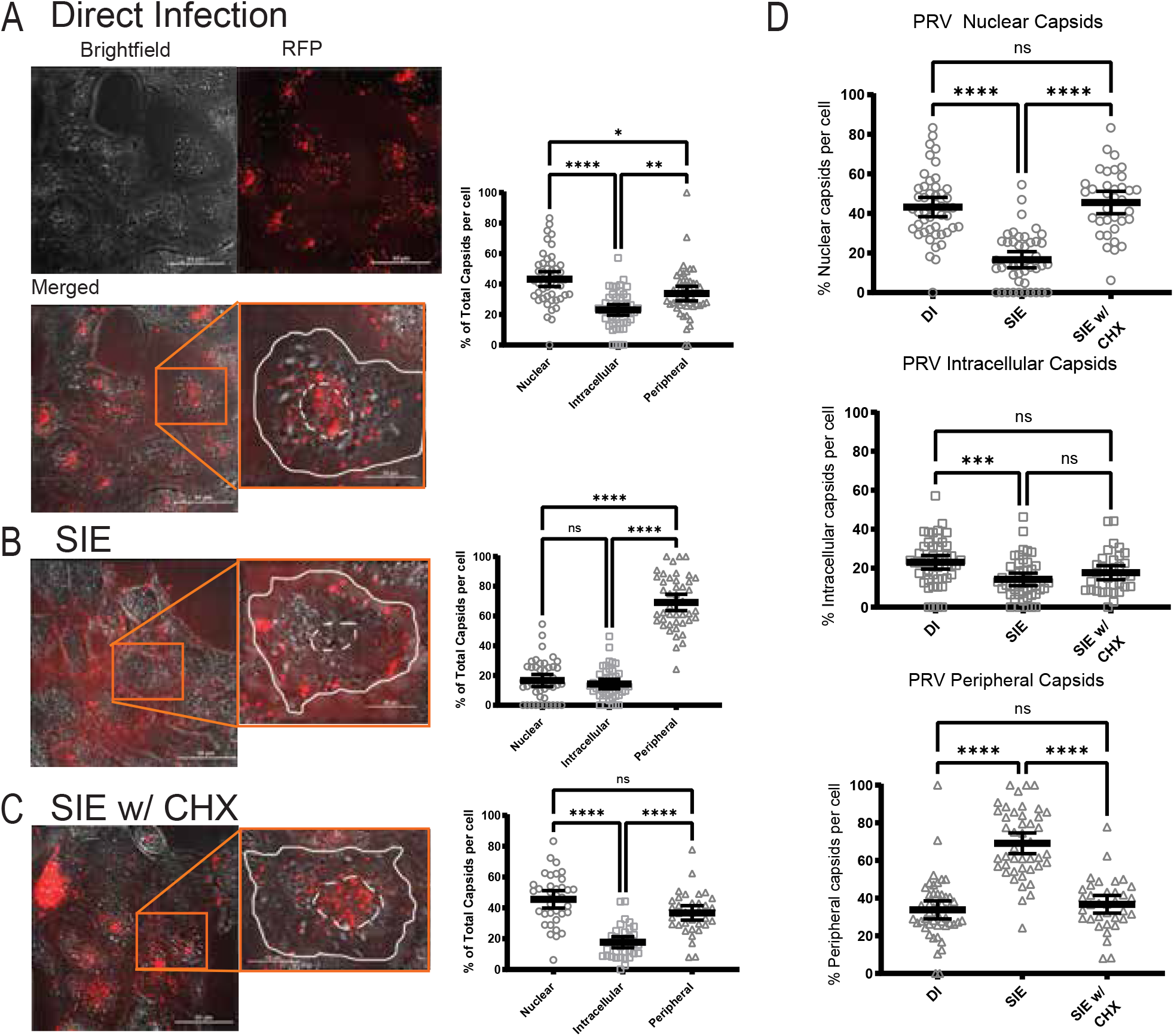
PRV SIE exhibits a strong reduction in nuclear associated capsids. PK15 cells under indicated conditions. Depicted channels are brightfield, red fluorescence channel, and merged. Inset shows magnified image of a representative single cell from the field. The white dotted line corresponds to the nuclear region of the cell, while a solid line depicts the periphery of the cell. For each condition, a minimum of 3 fields were analyzed and depicted as percent of total capsids per cell for each category. Quantitative results from single-cell analysis were statistically compared by One-Way ANOVA analysis. A) Direct infection with PRV 180 (MOI 100). B) SIE infection with primary infection of PRV Becker (MOI 100) with secondary infection of PRV 180. C) SIE with CHX treatment. CHX treatment occurred between infection of primary and secondary. D) Dunn’s multiple comparison test (One Way ANOVA) comparing capsid localization between conditions. * represents a p-value of 0.0129. ** represents a p-value of 0.0016. *** represents a p-value of 0.0004. **** represents a p-value of <0.0001.

**Figure 5.**
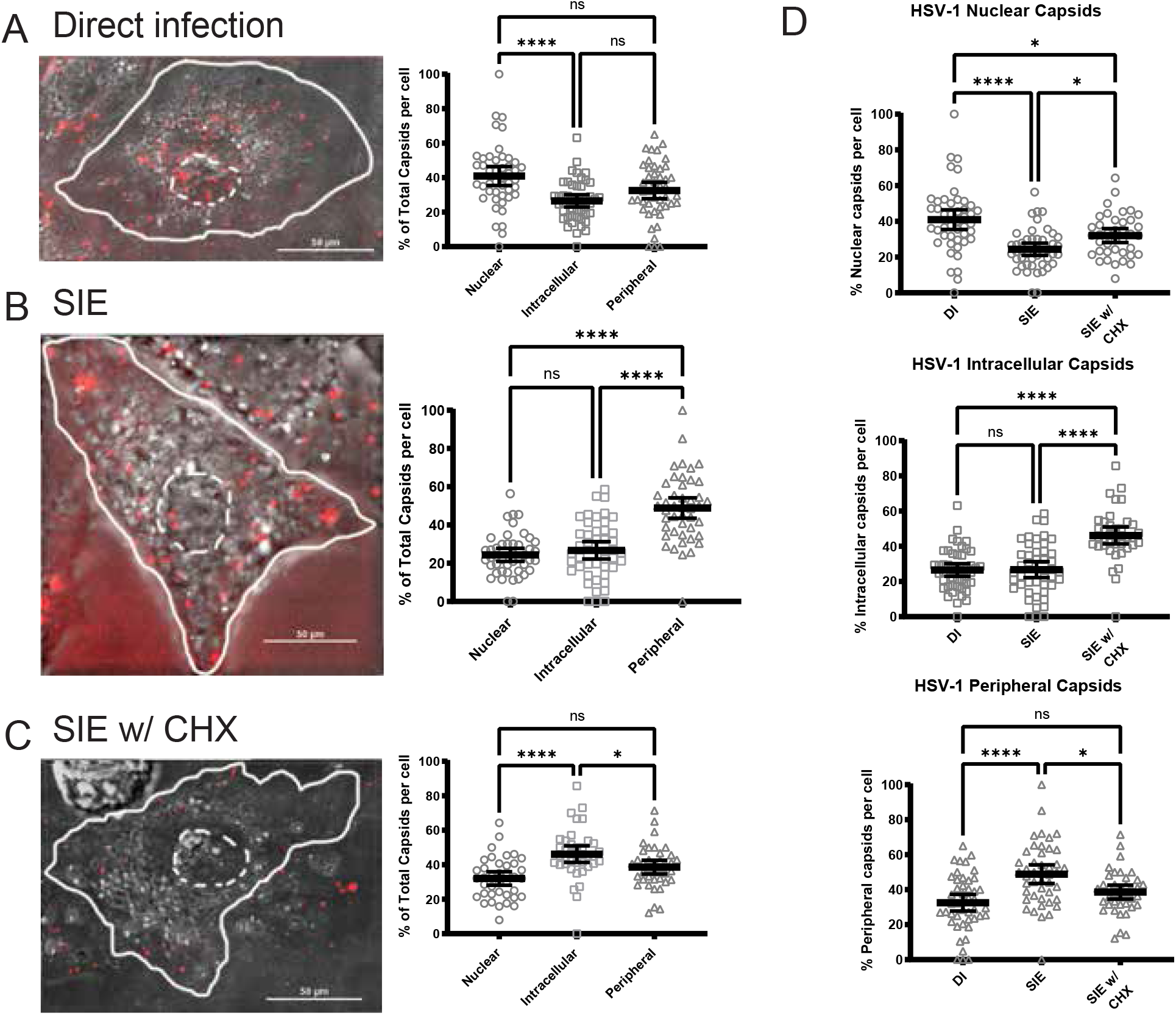
HSV SIE exhibits a modest reduction in nuclear capsids. Representative images of an infected cell with capsid distributions from three conditions of infection. The white dotted line indicates the nuclear region of the cell, while a solid line depicts the periphery of the cell. For each condition, a minimum of 3 fields were analyzed and depicted as percent of total capsids per cell for each category. Quantitative results from single-cell analysis were statistically compared by One-Way ANOVA analysis. A) Direct infection with HSV OK14 (MOI 100). B) SIE infection with primary infection of HSV-1 17 (MOI 100) and secondary infection of HSV OK14. C) SIE with CHX treatment. CHX treatment occurred between infection of primary and secondary. D) Dunn’s multiple comparison test (One Way ANOVA) comparing capsid localization between conditions. * represents a p-value of 0.0264. **** represents a p-value of <0.0001.

**Figure 6.**
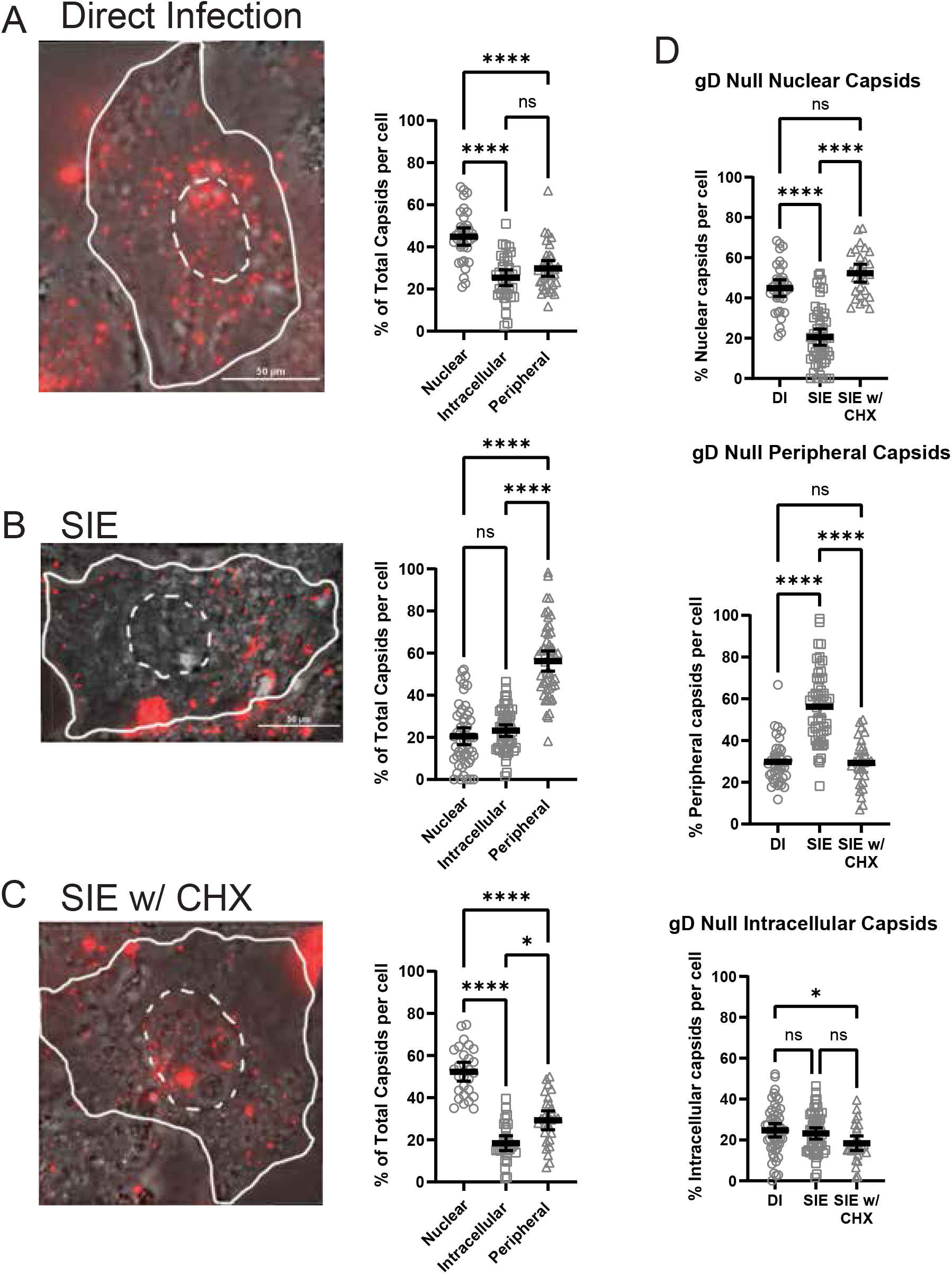
SIE during gD-null infection also reduces nuclear-associated capsids. Representative images of capsid distribution under three conditions of infection. The dotted white line indicates the nuclear region of the cell, while a solid line depicts the periphery of the cell. For each condition, a minimum of 3 fields were analyzed and depicted as percent of total capsids per cell for each category. Quantitative results from single-cell analysis were statistically compared by One-Way ANOVA analysis. A) Direct infection with PRV 180 (MOI 100). B) SIE infection with primary infection of PRV GS442 (gD null) with secondary infection of PRV 180. C) SIE infection with CHX treatment. CHX treatment occurred between infection of primary and secondary. D) Dunn’s multiple comparison Test (One Way ANOVA) comparing capsid localization between conditions. * represents a p-value of 0.0292. **** represents a p-value of <0.0001.

The counts and categorizations of fluorescent capsid assemblies for PRV are presented in Figure 4. We were able to quantify an average of 45 virions per cell, with a total of 35 to 48 cells within each condition. Plotted are the numbers of capsids categorized for each condition out of the total observed capsids on a per cell basis. We also compared capsid localization across different conditions to determine the statistical significance between infection conditions.

Comparisons between capsid distributions by condition reveal that there was a strong statistical difference between SIE and direct infection (DI) for all capsid categorizations (Figure 4a-c). For the DI condition, an average of 47% of cell-associated capsids were localized near the nucleus, while 35% were localized at the periphery. For the condition of SIE, only 14% of capsids localized near the nucleus, while 72% of total capsids were at or near the cell periphery. For the condition of SIE with CHX treatment, capsid distributions more closely matched DI conditions, with 47% of capsids at the nucleus and 37% were on the periphery of the cell. It is important to note that the values are averaged across multiple cells observed for each condition. As a result, most average values under-represent the most observed state of capsid distribution. For SIE, many cells contain no nuclear capsids and have 100% of capsids retained at the periphery. Comparisons of capsid distribution between SIE and DI revealed that there was a significant difference between the nuclear and peripheral distribution of capsids but not any statistical difference for capsids localized to the intracellular region of the cell. The overall reduction of nuclear and intracellular capsids and an accumulation of peripheral capsids during SIE correlates with our live fluorescent microscopy observations. Together, the differences in capsid distribution indicate that PRV-induced SIE inhibits entry.

### HSV-1 SIE does not alter intracellular capsid accumulation

To compare how SIE alters secondary capsid uptake between different alphaherpesviruses, we performed a parallel experiment with HSV-1 expressing an mRFP fusion to VP26 (HSV-1 OK14) under similar conditions. We quantified an average of 40 capsids per cell, analyzing between 39 and 46 cells per condition. The direct infection condition averaged 44% of total capsids localized to the nuclear region, while 27% of total capsids were localized to the periphery of the cell. The condition of SIE averaged 23% of total capsids within the nuclear region, while approximately 52% of virions remained on the periphery of the cell. The condition of SIE with CHX treatment was more intermediate between conditions, with approximately 33% localization of capsids to the nuclear region of the cell, while 40% of red puncta localized to the periphery of the cell.

Results from the quantification of mRFP-labeled HSV-1 capsids revealed a surprising difference of capsid distribution compared to PRV. As shown in Figure 5d, there was a reduction of nuclear capsids and an increase in peripheral capsids between HSV SIE and HSV DI. However, there was no statistical difference between their intracellular capsid count. This differs from PRV, where infection during SIE exhibited an overall reduction of capsid internalization. Moreover, treatment with CHX in HSV-1 SIE elicited a slight decrease in capsid internalization, a strong increase in intracellular capsids, and no significant increase in peripheral capsids. These differences in capsid distribution seem to indicate that HSV does not interfere with superinfecting virion entry to the same extent as PRV.

### PRV gD-expression is not necessary to reduce capsid trafficking

We previously determined that SIE for PRV occurred independently of gD expression (4). However, it does not eliminate the possibility that gD-expression could be interfering with the entry of the superinfecting virion. To test if gD expression is necessary for the observed reduction in capsid entry during PRV SIE, a gD-null virus (PRV GS442) was utilized as a primary inoculum.

Using the same conditions as already defined in figures 4, an average number of 58 fluorescent capsids were quantified for 30 to 55 cells for each condition. Although this is a greater average count per cell in comparison to the prior analyses, aggregates of fluorescent capsids were more extensive during superinfection of gD-null infected cells. Cells with excessive capsid aggregation were excluded from quantification with our analysis focusing on cells with individual capsid assemblies. The direct infection condition averaged 44% of total capsids localized to the nuclear region, while 30% of total capsids were localized to the periphery of the cell. The condition of SIE averaged 22% of total capsids within the nuclear region, while approximately 55% of capsids remained on the periphery of the cell. The condition of SIE with CHX treatment demonstrated a similar profile to direct infection with approximately 55% localization to the nuclear region of the cell, while 28% of red puncta localized to the periphery of the cell.

SIE with gD-null primary inoculation was very similar to our previous observation of decreased capsid association with the nucleus and accumulation of capsids at the periphery. Comparing the capsid distributions demonstrated a significant reduction (approximately 50%) for nuclear internalization between DI and SIE with no difference between DI and SIE with CHX treatment. There is an increase of capsids on the periphery of the cell for SIE only with no differences between DI and SIE with CHX treatment. The only distinct difference between wild-type SIE and gD null SIE is the percent of intracellular capsids not associated with the nucleus. This lack of difference in internalized capsids may be related to the accumulation of fluorescent capsid assemblies at the periphery or to changes in secondary virion fusion compared to wild-type PRV primary infection. Despite the subtle changes in capsid uptake, the reduction in nuclear capsid association following gD-null primary infection is consistent with a mechanism inhibiting virion entry that is independent of gD expression.

## DISCUSSION

This study probed the effect of glycoprotein-D independent SIE during secondary entry of the alphaherpesviruses PRV and HSV-1. We observed that the MOI for both the primary and secondary inoculum played a significant role in the establishment and extent of exclusion. A higher MOI secondary inoculum was able to overcome SIE, while lower MOIs of secondary challenge are more efficiently excluded. One possible explanation for these observations is that higher MOI primary infections exhibit expedited viral gene expression and implementation of SIE. For example, an MOI of 10 for the initial virus completely failed to exclude a superinfecting MOI of 25, while an initial MOI of 25 required a superinfecting MOI of 100 to achieve a similar expression of SIE. Additionally, conditions with a higher initial MOI are more capable of excluding the secondary virus (as shown in the columns when comparing conditions like MOI 50 and 100 with a lower secondary viral infection). This suggests that the kinetics of viral replication and genome production likely play roles in regulating implementation of SIE. The importance of primary viral genome number and subsequent protein production was further confirmed using CHX to inhibit protein production by the primary viral infection. High or low inoculation with similar MOIs and CHX treatment alleviated SIE. This indicates that it is not just the presence of a high inoculating virus, but also its amount relative to the secondary virus.

Although concurrent results indicate viral factors can influence the extent of SIE, nonviral factors could not be excluded. To this end, we evaluated surface expression of nectin-1 during infection. Although there is a drop in nectin-1 following initial infection at 0 hours post infection, the level of surface nectin-1 is similar when treated with CHX. Since CHX has been previously shown to alleviate SIE, the limited nectin-1 present on the surface is still capable of mediating superinfection. This observation strongly suggests that interference with cell surface presentation of entry receptors is not important for the mechanism of alphaherpesvirus SIE.

To address the step(s) involved with gD-independent SIE, we visualized capsid entry with live fluorescent microscopy. Dual-labeled virions allowed the observation of distinct fluorescent profiles for virion distribution between SIE and direct infection conditions. Accumulation of dual fluorescent PRV virions at the plasma membrane of cells during SIE suggests an inhibition of virion fusion and confirms prevention of entry. This is further supported by the quantitative analysis of fluorescent capsids which revealed strong reductions in nuclear association following entry. These results raise the possibility that gD-independent SIE might inhibit virion entry or subsequent trafficking.

Initially, it was expected that the mechanism for SIE would be similar across alphaherpesviruses. Experiments from Criddle et. al. demonstrated that HSV-1 and PRV SIE shared similar attributes to inhibit secondary inoculum associated FP expression. However, the results from this study indicate that this is not the case. PRV-mediated SIE results in reduced capsid internalization, with most accumulating on the periphery. HSV-1 SIE, however, doesn’t exhibit reduced capsid internalization while still resulting in accumulation on the periphery. This result indicates that HSV-1 SIE might employ a combination of post-entry factors to suppress expression of a superinfecting virus, while PRV SIE likely targets only virion fusion (Figure 7). This difference strongly suggests there are multiple mechanisms employed by alphaherpesviruses to suppress superinfection. The single-cell analysis of capsid distribution is one potential means for continued study into the differences of SIE between related viruses.

**Figure 7.**
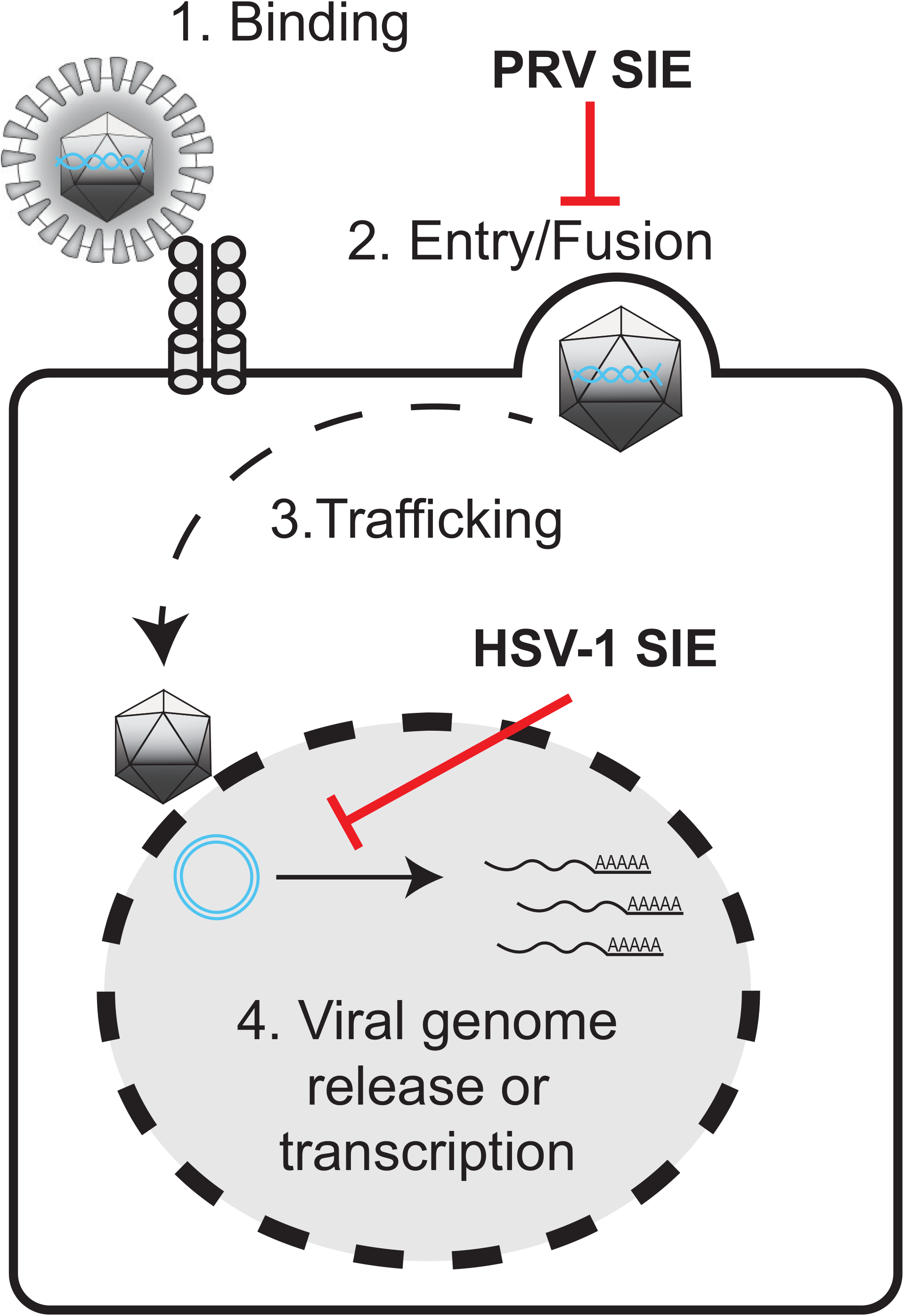
Model of SIE inhibition of alphaherpesviruses. A schematic representation of the proposed points of inhibition of secondary virion infection during SIE. PRV SIE suppresses entry/fusion, while HSV-1 possibly inhibits viral genome release or transcription.

Taken together, work from this study provides new insights into the properties of alphaherpesvirus SIE. This work supports our previous finding that SIE is a virally induced, gD-independent phenotype but differs in the mechanism of exclusion between PRV and HSV-1. PRV mediated SIE appears to inhibit entry of secondary virions in a gD-independent manner, while HSV-1 SIE impacts a post-entry step. Further work will further characterize the differences between PRV and HSV-1 SIE, as well identify the viral mechanism(s) which mediate exclusion. Most likely, this will address the kinetics of viral protein between primary and secondary challenge of viruses. This data about gD-independent SIE will provide valuable insight regarding alphaherpesvirus population dynamics, disease kinetics for neuronal pathologies, and development of antiviral therapies.

## ACKNOWLEDGEMENTS

The authors appreciate the sharing of reagents by the Smith and Enquist labs. We also thank Kyle Hain for providing critical reading and commentary of the manuscript. This work was supported by National Institutes of Health R21 Exploratory/Development Award Grants [R21AI139935] and [R21AI146952].

## Supplemental Movie legends

**Supplemental Movie 1. Live fluorescent microscopy of direct infection.** Movie of direct infection with PRV 137. Images taken every 2.5 minutes for 1 hour. Playback speed is 5 frames per second. Scale is equal to 10 μm.

**Supplemental Movie 2. Live fluorescent microscopy of SIE.** Movie of SIE with secondary inoculation of PRV 137, primary with nonfluorescent PRV Becker. Images taken every 2.5 minutes for 1 hour. Playback speed is 5 frames per second. Scale is equal to 10 μm.

